# Mechanical mismatch between Ras transformed and untransformed epithelial cells

**DOI:** 10.1101/152942

**Authors:** Corinne Gullekson, Gheorghe Cojoc, Mirjam Schürmann, Jochen Guck, Andrew Pelling

## Abstract

The organization of the actin cytoskeleton plays a key role in regulating cell mechanics. It is fundamentally altered during transformation, affecting how cells interact with their environment. We investigated mechanical properties of cells expressing constitutively active, oncogenic Ras (Ras^V12^) in adherent and suspended states. To do this, we utilized atomic force microscopy and a microfluidic optical stretcher. We found that adherent cells stiffen and suspended cells soften with the expression of constitutively active Ras. The effect on adherent cells was reversed when contractility was inhibited with the ROCK inhibitor Y-27632, resulting in softer Ras^V12^ cells. Our findings suggest that increased ROCK activity as a result of Ras has opposite effects on suspended and adhered cells. Our results also establish the importance of the activation of ROCK by Ras and its effect on cell mechanics.

## INTRODUCTION

The actin cytoskeleton is largely responsible for the mechanical properties of the cell. A myriad of actin-binding proteins regulate actin filament assembly and the organization of actin filament networks ^1^. Actin filaments are organized into the cortex, a thin cross-linked mesh lying immediately beneath the plasma membrane, and fibers, contractile actin bundles that are anchored by cell adhesions ^1–3^. The actin cytoskeleton is fundamentally different in adherent and suspended cells. It has been shown that that the mechanics of cells change after they detach from substrates ^4,5^. Adherent cells have anchoring points to the substrate, called focal adhesions, and actin stress fibers span through the cell from one focal adhesion point to another. Attachment to a substrate is thus a prerequisite for the formation of stress fibers and they are not observed in non-adherent cells. In epithelial monolayers, there is also a band of cell-cell adhesions, called adherens junctions, that encircles the cell ^6^. These attachments act as anchors for a circumferential belt of actin fibers inside the cell ^7,8^. The molecular motor, myosin, can interact and modify the cytoskeleton altering the mechanical properties of the cell. Myosin can generate tension along actin fibers^9^ and pre-stress the actin cortex ^10^. Cell stiffness increases with the level of tensile stress or pre-stress in the cell ^11^. Seemingly due to decreasing pre-stress in the actin network, it has been shown that both the decrease in stress fibers on compliant substrates and the inhibition of myosin on substrates correlate with the softening of cells ^12,13^. However in suspension, the inhibition of myosin correlates to the stiffening of cells ^14^. Similarly it has been shown that the formation of adherens junctions coincide with an increase in the apparent stiffness of reforming cell monolayers ^15^.

Cell transformation can arise from the activation of oncoproteins. A common oncoprotein is Ras, which controls a variety of cellular signaling pathways responsible for growth, migration, adhesion, cytoskeletal integrity, survival and differentiation ^16^. Mutations that permanently activate Ras are found in approximately 20% of cancers ^17^. For this reason, there is great interest in developing tumor therapies targeting Ras and Ras effector pathways ^18^. One pathway of particular importance for cell mechanics is the one activating contraction through Rho, Rho-associated protein kinase (ROCK) and myosin light-chain kinase (MLCK). Ras activity is associated with increased Rho activity ^19–22^. Rho activates ROCK, which regulates actin filament organization and actomyosin contractility ^23,24^. It has been observed that phosphorylation of myosin light chain is enhanced in cells with permanently active Ras^V12 25^. The over-activation of this pathway should have different effects on adherent and suspended cells due to their difference in cytoskeletal structure.

During early stages of tumor progression in an epithelial monolayer, cells of different mechanical stiffness are in contact with one another. The mechanical mismatch between transformed and non-transformed cells could be an important driver of cancer progression. Typically, it has been shown that cancerous cells are softer than non-transformed cells and that highly aggressive cancerous cells are softer than less aggressive cancerous cells ^26–29^. This can be attributed to the cytoskeleton devolving from a rather ordered and rigid structure to a more irregular and compliant state ^27^. It has also been shown using a human breast cell line with conditional Src induction, that cells undergo a stiffening state prior to acquiring malignant features ^30^. Metastatic cancer cells are also in different states of attachment during their journey from one site to another. During the initial stages the cell is in a monolayer with cell-cell and cell-substrate adhesions. When the cell is in the blood stream or lymphatic system, it is in a suspended state. It is thus important to measure cell mechanics in different degrees of attachment.

A popular method of measuring mechanical properties of adherent cells is atomic force microscopy (AFM), where the surface of the cell is indented by a probe consisting of a cantilever with a sharp tip ^31,32^. This is generally preformed on cells adhered to stiff tissue culture dishes made of plastic or glass. An optical stretcher (OS) is an orthogonal method for measuring the mechanical properties of cells in suspension where no adhesion is needed ^33^. The OS stretches the cell with optically induced stresses on the cell surface ^34^. The distribution of stress applied to the cell is broad and continuous. This global stress application is in contrast to the local deformations achieved with atomic force microscopy that can encounter artifacts due to inhomogeneous cell morphology. However, these techniques both use small deformations to measure the cortical tension of the cell. Moreover, there is a major difference in the attachment states of the cells measured by these two techniques. These two techniques allow us to measure the two extremes of cellular attachment, allowing us to model the different attachment of cells and determine their effect on cell mechanics.

In this study, we investigated the effects of Ras on the mechanical properties of attached and suspended epithelial cells. A cell line that produces constitutively active Ras in MDCK cells was utilized for this purpose ^25^. Y-27632 was used to inhibit actomyosin contractility; it inhibits ROCK, which is downstream of Ras ^35^. Our results show that constitutively active Ras increased the stiffness of cells in monolayer and decreased the stiffness of suspended cells. When ROCK was inhibited with Y-27632, the Ras^V12^ cells were softer in monolayer than normal cells. When in suspension, the inhibition of ROCK had an opposite effect and stiffened the Ras^V12^ cells. Our work provides evidence that Ras plays a role in stiffening cells in monolayers on stiff substrates and softening cells in suspension. This demonstrates that Ras has opposite effects on the stiffness of cells in attached and suspended states.

## METHODS

### Cell Culture

Madin-Darby Canine Kidney (MDCK) epithelial cells and Ras^V12^, a stable MDCK cell line expressing GFP-Ras^V12^, in a tetracycline-inducible manner were cultured in DMEM with 10% fetal bovine serum (FBS), 50mg/ml streptomycin and 50U/ml penicillin antibiotics (all from Hyclone Laboratories Inc.). The Ras^V12^ line was a kind gift of Yasuyuki Fujita (Hokkaido Univ) described here ^25^. Cells were cultured at 37°C in a 5% CO2 incubator on 100mm tissue culture dishes (Corning). Samples were either grown in mono-culture or in co-culture with a ratio of 1:10 Ras^V12^ to wild type (WT) MDCK cells to create monolayers. The Ras^V12^ cells were activated with tetracycline at a concentration of 2μg/ml and experiments were performed 1 day later. The inhibitor Y-27632 was used at a concentration of 10μM in a 1 day treatment.

### Imaging

Stained images were acquired on a TiE A1-R laser scanning confocal microscope (LSCM) (Nikon) with a 60X objective. Images were acquired with a standard LSCM configuration with appropriate laser lines and filter blocks. Cells were fixed with 3.5% paraformaldehyde and permeabilized with Triton X-100 at 37°C. DNA was labeled with DAPI (Invitrogen). Actin was stained with Phalloidin Alexa Fluor 546 (Invitrogen). Actin signal intensities at cell boundaries were measured in ImageJ with line segments placed over cell margins in images of MDCK WT:Ras^V12^ co-cultures. The boundaries were either between two Ras^V12^ cells (Ras^V12^:Ras^V12^, mean of 11 boundaries per image), a Ras^V12^ cell and a neighboring WT cell (Ras^V12^:WT, mean of 28 boundaries per image) or the neighboring WT cell and another WT cell (WT:WT, mean of 41 boundaries per image). The average intensity over each boundary type was calculated for each image (5 images per condition) and divided by the average intensity of the WT:WT boarder.

### Atomic Force Microscopy

A NanoWizard II (JPK Instruments) AFM was used with PNP-TR (nanoworld) cantilevers (68±1pN/nm). The AFM was mounted on a TiE A1-R laser scanning confocal microscope (Nikon). All experiments were performed at 37°C, using a temperature-controlled AFM stage (JPK). Cells were grown on glass bottomed dished coated with collagen. Force-curves were measured over the center of cell nuclei and fit to the Hertz model for a conical tip (200nm indentations) in order to derive the local Young’s modulus (PUNIAS Software). At least 23 cells were probed for each condition.

### Digital Holographic Microscope

Wild type and Ras^V12^ cells were trypsinized and resuspended in phosphate buffered saline (PBS). The PBS has a known refractive index of 1.335 determined by a refractometer (2WAJ, Arcarda GmbH). The cells in suspension were put on a glass cover slip coated with poly-l-lysine that was used to produce a charge-based attachment for the cells. Images of the cells were taken with a Digital Holographic Microscope a short time period after settling, so that they retained the spherical shape from the suspended state. The setup and image analysis is described in detail here ^36^. In short, the hologram image was used to determine the quantitative phase shift of light induced by the sample. A circle was fit to the contour of the cell and the cell was assumed to be a sphere to determine the height of the sample for each pixel. This was used to get the average refractive index of the cell.

### Optical Stretcher

The working principle of a microfluidic optical stretcher has been described in detail here ^27,37^. The device is a dual beam laser trap capable of trapping and deforming cells through optically induced stress acting on the cell surface. The microfluidic flow chamber consists of a square glass capillary, aligned perpendicular to two single mode optical fibers, which delivers the cells in suspension. Cells were stretched within one hour of being detached from a culture dish with trypsin. Single cells were trapped with the laser at low power and held for 2 seconds to stabilize. Subsequently, the cells were stretched at high power for 4 seconds. This might introduce heating effects that can impact cell mechanics ^38^. The strain (*γ*) at each time point was calculated using change in length of the cell along the axis the stress was applied.

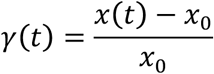

The compliance (*J*) was calculated by dividing the strain by the peak stress on the cell (*σ*_0_) and a geometric factor (*F_g_*) described here ^39^.

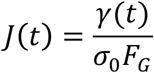

For each cell type under different conditions, the number of selected cells per OS experiment was n≥30. The creep compliance data are depicted as mean±SE. All measurements were performed with the cells suspended in phosphate buffered saline (PBS) at room temperature. All plotting, statistical analysis, and curve fitting were performed using MATLAB.

## RESULTS

Ras^V12^ cells were examined in different states of attachment. We created a high state of attachment by seeding Ras^V12^ cells in in co-culture with WT cells on collagen-coated glass. The mixture of cells was cultured in the absence of tetracycline until they formed an epithelial monolayer. Tetracycline was then added in the culture medium to induce Ras^V12^ expression, simulating the transformation of a cell in monolayer surrounded by untransformed cells. The cells were examined 24 hours after transformation to examine cell stiffness by AFM as well as basal protrusion, apical extrusion and actin signal. To remove cell-cell interactions single cells loosely attached to glass were also probed by AFM. Suspended cells were allowed to attach for 30min before having their stiffness measured. A fully suspended state with no cell-cell or cell-substrate attachments was also examined with a microfluidic optical stretcher.

### Ras stiffens cells in Monolayer

To investigate the effects of constitutively active Ras on attached monolayers, co-cultures of Ras^V12^ and WT were studied (Fig. 1A). Many Ras^V12^ cells were observed to be extruding apically out of the monolayer or protruding below the monolayer (Fig. 1B). When stained for actin, the signal was more intense in the margins of the Ras^V12^ cells both at the interfaces between two Ras^V12^ cells (Ras^V12^:Ras^V12^) and interfaces between Ras^V12^ and WT cells (Ras^V12^:WT) than the interfaces between two WT cells (WT:WT) (p<0.05) (Fig 2A). The actin signal was also more intense at the Ras^V12^:Ras^V12^ interfaces than the Ras^V12^:WT interfaces (p<0.05). More of the Ras^V12^ cells were protruding basal than were apically extruding (Fig 2B). This was consistent with the observations by Hogan et al ^25^. The cells in monolayer were probed with an AFM to measure stiffness (Fig. 2C). Force curves taken over the cell nuclei were fit to determine apparent Young’s moduli of the cortical membrane. The Ras^V12^ cells were stiffer than WT cells (p<0.05).

**Figure 1:**
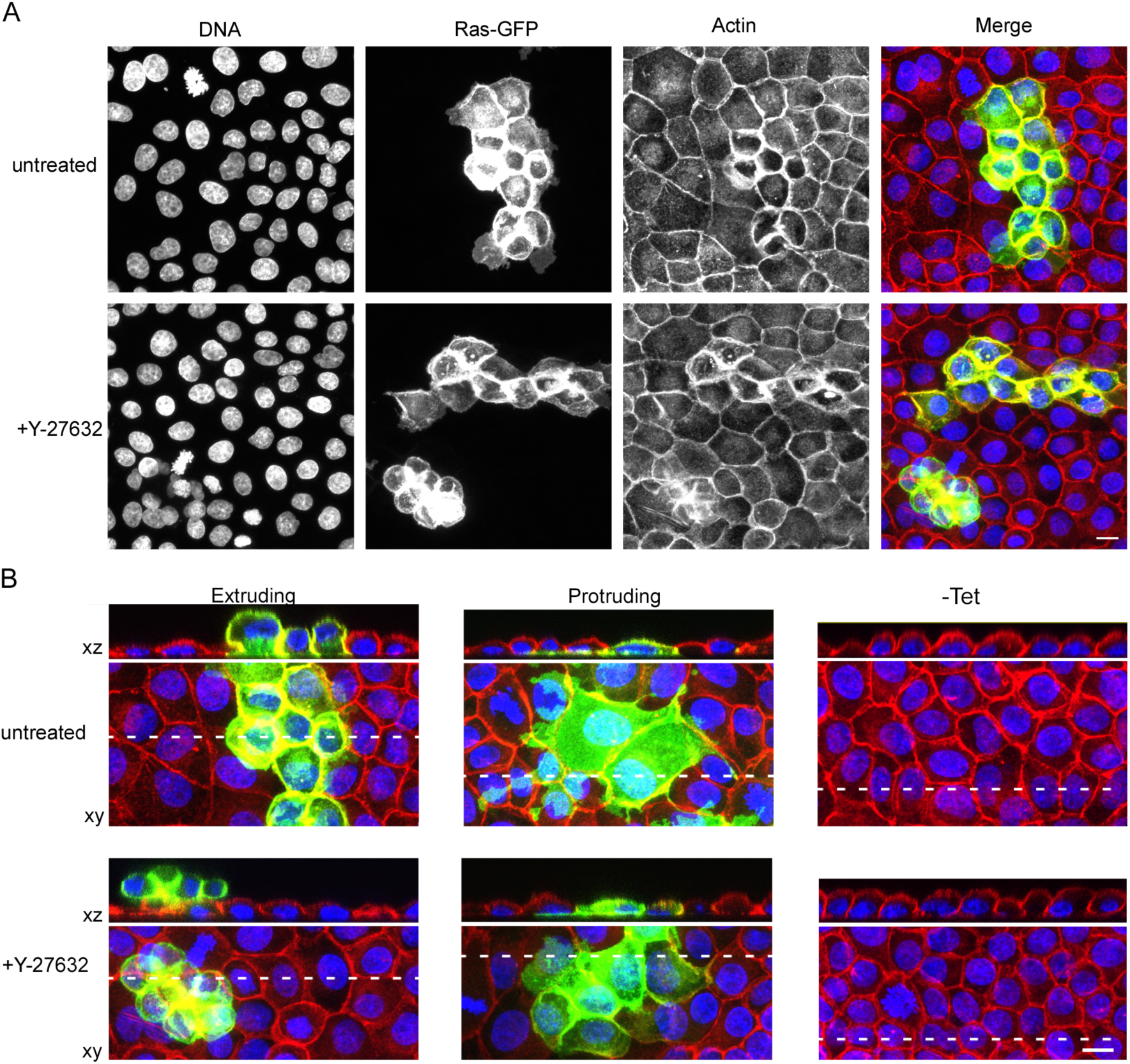
Morphology of Ras transformed cells in monolayer with WT cells. A) Maximum projections of confocal images of Ras^V12^ (green) and WT cells in monolayer with tet and with/without Y-27632 stained for DNA (blue) and actin (red) B) Orthogonal views and their corresponding maximum projections of Ras^V12^ and WT in monolayer, with/without tet, with/without Y-27632. White dashed lines indicate the positions of the xz planes used for the orthogonal views. Scale bars=10*μ*m

**Figure 2:**
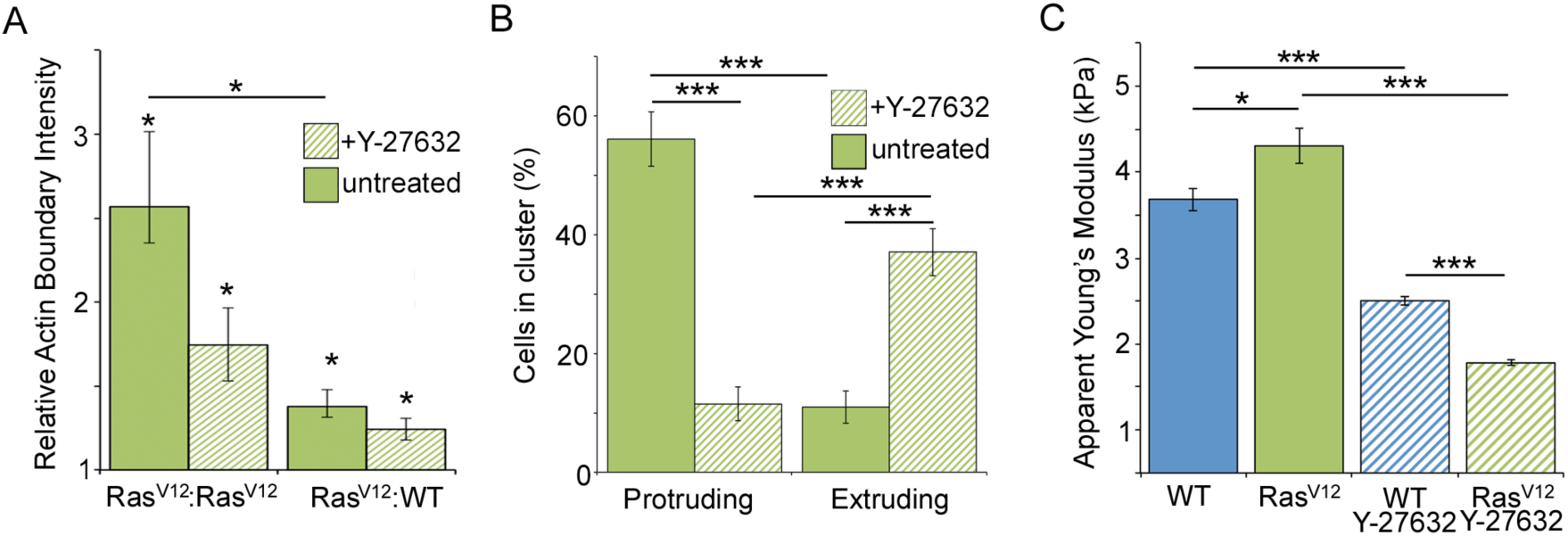
Properties of Ras transformed cells in a monolayer with WT cells. A) Average intensity of actin signal around the boundaries of cells either at the interface of Ras^V12^: Ras^V12^ cells and Ras^V12^:WT cells with and without Y-27632 relative to the intensity at the WT:WT interface. Each case is significantly greater than 1. B) Percent of Ras^V12^ cells in a cluster that are protruding or extruding with or without Y-27632. C) The AFM apparent Young’s moduli of WT cells and Ras^V12^ cells in co-culture, with and without the addition of Y-27632. Bars represent standard error of the mean, *p<0.05 ***p<0.001.

In order to investigate if the stiffer Ras cells were a result of increased ROCK activation/contraction, the inhibitor Y-27632 was employed. The monolayers were inhibited for 24h before testing, simultaneously with the tetracycline treatment. There was still a larger actin signal around the Ras^V12^ cells than at WT: WT interfaces (p<0.05) (Fig 2A). Apical extrusion and basal protrusion of Ras^V12^ cells were still observed with this treatment, however extrusion was less frequent and protrusions were more frequent (p<0.001)(Fig 2B). This decrease in extrusion was also observed in MDCK cells overexpressing Ras with the co-expression of dominant-negative ROCK ^25^ and in MDCK cells overexpressing Src with Y-27632^40^. As expected, for all samples the Y-27632 treated cells were softer than the untreated sample (p<0.001) (Fig 2C). This is consistent with other AFM studies using Y-27632 on NIH3T3 fibroblast ^41^, MDCK-II epithelial ^15^ and MDA-MB-231 metastatic breast cancer ^42^ cell lines. The wild type cells were stiffer than the Ras^V12^ cells (p<0.001) when treated with Y-27632, which is the opposite of what was found in the untreated condition. This demonstrates that the ROCK inhibitor softens the cells in monolayer. The ROCK inhibitor had a greater effect on the Ras^V12^ cells, making the Ras^V12^ cells softer than the WT cells. Since the Ras^V12^ cells were softer than WT cells when ROCK was inhibited, it is likely that there are other Ras pathways affecting cytoskeletal organization, attributing for the softening of the cell. The difference in relative stiffness may account for the differences in extrusion direction in the Y27632 and untreated cases.

### Ras softens loosely attached cells

Based on the initial AFM findings we sought to examine the influence of cell-cell adhesion on the physical properties of the cell. Therefore, we carried out the following experiment where single cells were loosely attached to dishes. Cells were removed from a monolayer with trypsin. They were then resuspended in new media and transferred to a fresh glass bottomed dish. After 30min in the new dish, the cells attached to the glass allowing the cells to be probed by AFM (Fig 3A). This allowed us to probe cells by AFM while in a state with no cell-cell adhesions. With this method we can see if cell-cell adhesions are needed for the stiffening effect of Ras. The Ras^V12^ cells were softer than WT cells (p<0.001) (Fig 3B). This supports the theory that the stiffer Ras^V12^ cells in monolayer were the result of increased actomyosin contractility in the circumferential actin belt.

**Figure 3:**
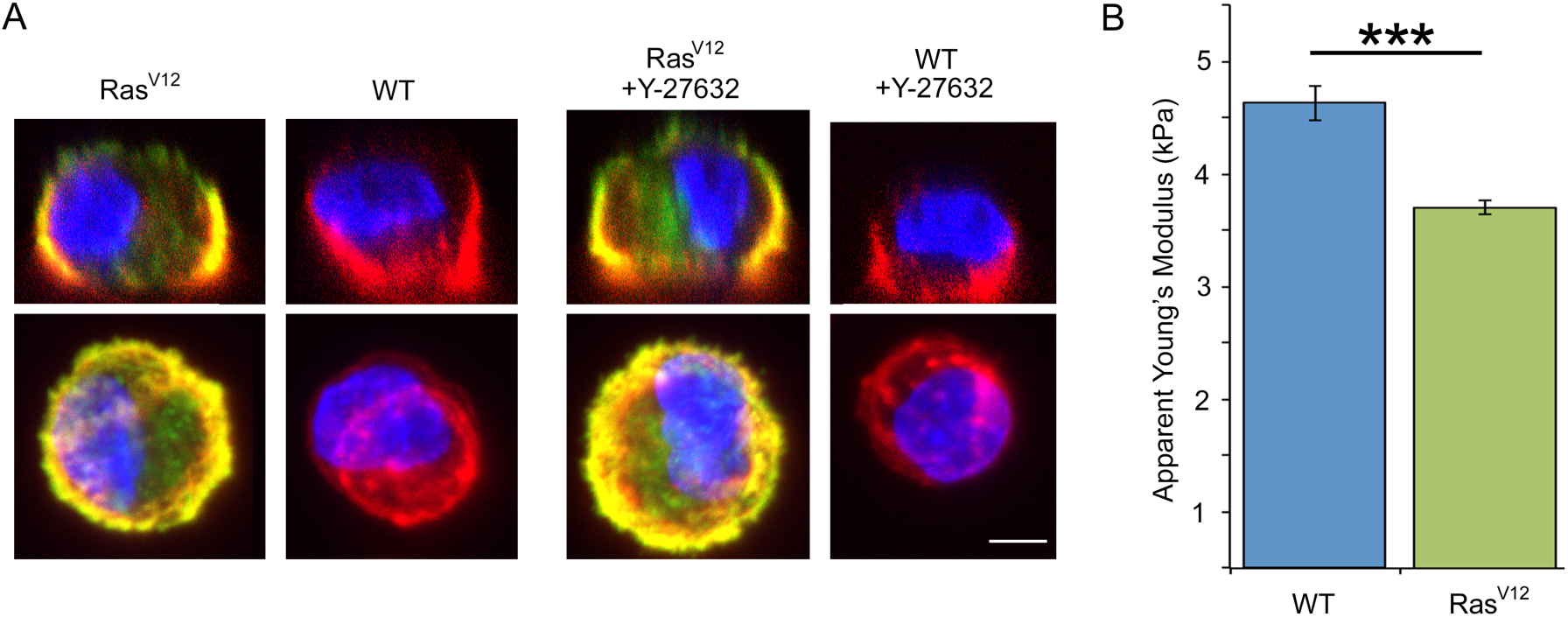
Influence of Ras on the apparent stiffness of cells loosely attached to glass probed by AFM. A) Orthogonal views of confocal images of loosely attached Ras^V12^ cells and WT cells with and without the addition of Y-27632. B) The AFM apparent stiffness of WT cells and Ras^V12^ cells grown in co-culture. Bars represent standard error of the mean, Scale bar=5*μ*m, ***p<0.001.

### Ras softens cells in suspension

Based on the contrasting results found with fully attached and loosely attached cells measured by AFM, we focused on investigating fully suspended cells. Cells detached from Ras^V12^ and WT monolayers were examined in an optical stretcher. In this sate there are no cell-cell adhesions or cell-substrate adhesions.

Along with other parameters, the refractive index of the cell is needed in order to calculate the stress it experiences. Refractive index is also related to mass density in biological material ^43,44^, and gives some insight into the structure of the cell. By the fitting to the cell contour during the analysis of the DHM images the cell radius is also determined. Cells taken from WT and Ras^V12^ monocultures were examined with this method. The Ras^V12^ cells were found to be larger than the WT cells (p<0.001) and had a smaller refractive index (p<0.001) (Fig 4). It has been found previously that refractive index decreases with increasing size in pancreas tumor cell types ^45^ and that activation of Ras increases cell size in the drosophila wing ^46^. Therefore it is not surprising to observe the same trend in the Ras^V12^ line. The correlation between cell size and refractive index may be explained by increased cell water content ^47^.

**Figure 4:**
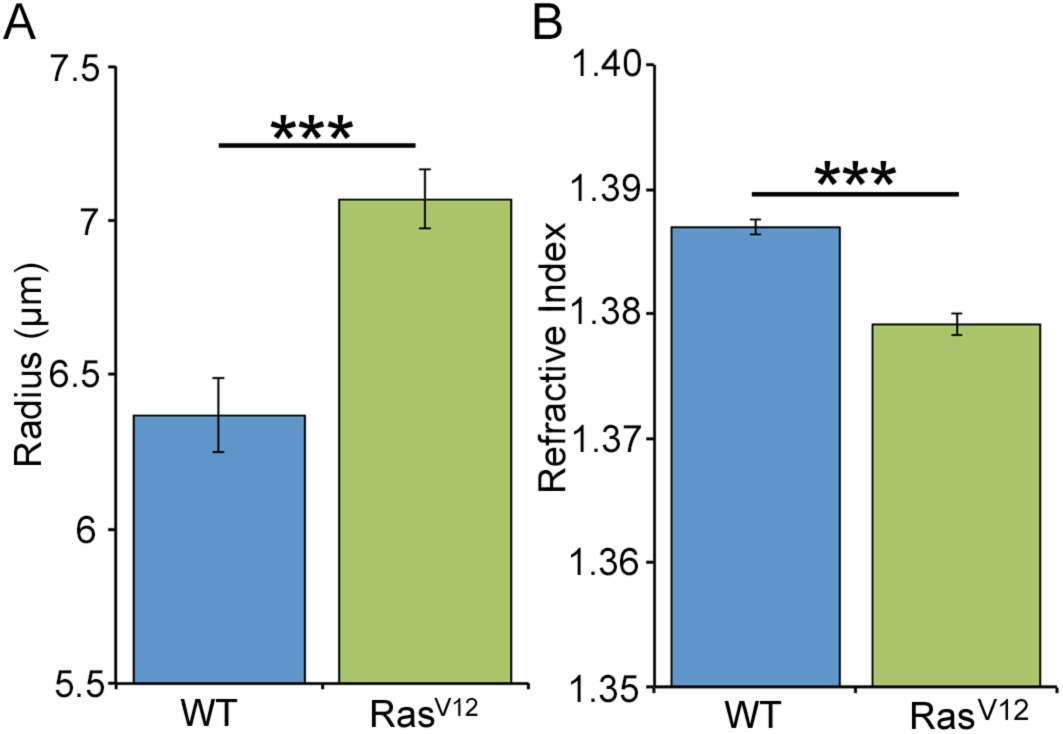
Effect of Ras on size and refractive index of cells measured with a digital holographic microscope. A) The radius of WT (n=52) and Ras^V12^ (n=54) cells. B) The refractive index of WT and Ras^V12^ cells. Bars represent standard error of the mean, ***p<0.001.

When measured in the optical stretcher, Ras^V12^ cells were found to have a higher compliance at the end of the 4 second stretching than WT (p<0.001) (Fig 5). This data demonstrates that the Ras^V12^ cells were more compliant than WT cells when in suspension. This is in contrast to what is seen in monolayers where the Ras cells had a higher effective Young’s Modulus. When treated with Y-27632 there was no significant difference of compliance at the end of stretching between the WT and Ras^V12^ cells (p>0.05). However, the addition of Y-27632 did lower the compliance of the Ras^V12^ (p<0.05) while having no significant effect on WT cells (p>0.05). This could be due to the increased ROCK/myosin light chain activity in the Ras cells. This is in contrast to what was seen in monolayers where the ROCK inhibition softened the cells. It has been found previously that inhibition of myosin decreases compliance of adherent cells brought into suspension ^14^.

**Figure 5:**
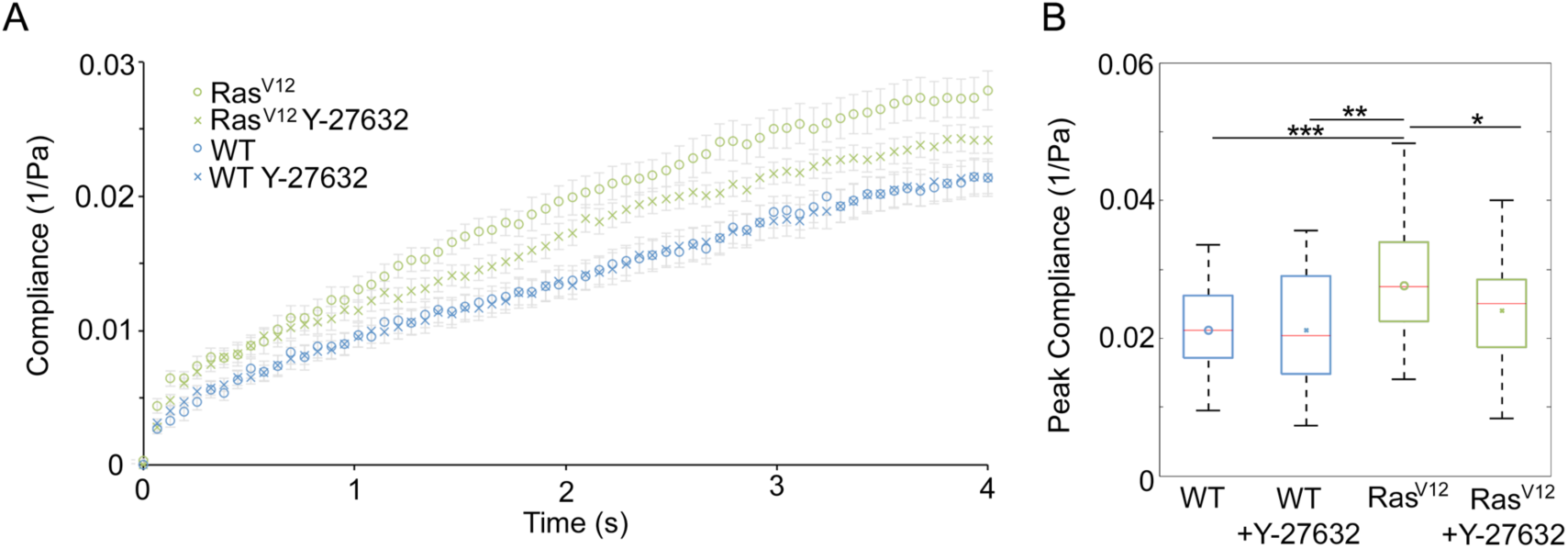
Influence of Ras on the compliance of cells in suspension probed by optical stretcher. A) The compliance curves of untreated and Y-27632 treated MDCK and Ras^V12^ cells. B) Box plots of the peak compliance (at 4 seconds) of each compliance curve. *p<0.05, **p<0.01, ***p<0.001.

## DISCUSSION

Ras has two major effects on the cell cytoskeleton; it alters cytoskeleton organization and contracts it. In cancer, the cytoskeleton is typically intrinsically softer due to its irregular structure ^27,48–50^. It has been shown that the amount of filamentous actin in malignantly transformed fibroblast cells is reduced by ~40% ^27^. However, contraction of cells by the activation of ROCK has a stiffening effect on adhered cells ^41,42,51^. This was reaffirmed with Y-27632 treated WT and Ras^V12^ cells being softer than their untreated counterparts when probed in monolayer by AFM. We found that Ras^V12^ cells were stiffer than WT cells when in the monolayer, but softer than the WT cells when ROCK activity was inhibited with Y-27632. We also found a greater actin signal in the cell margins of Ras^V12^ cells. The stiffing of Ras^V12^ cells in monolayer co-culture we observed is likely a result of increased pre-stress and cortical tension from the actomyosin contraction. This effect was likely removed with the ROCK inhibitor, showing the softening effects of the altered cytoskeleton organization effect of Ras. Adhesion on compliant substrates leads to less pronounced or even absent stress fibers ^52^. The formation of stress fibers could be an artifact of tissue culture, making this imperfect model of the *in vivo* environment.

When cell-cell interactions were removed by loosely attaching the cells, the Ras^V12^ cells were found to be softer than WT cells. When measured in the completely unattached state by applying stress optically, the Ras activated cells were softer highlighting the effect of altered cytoskeleton structure on cell stiffness. This demonstrates that cell-cell adhesion is necessary for the Ras^V12^ cells to be stiffer than WT cells. The suspended Ras^V12^ cells were also stiffer when ROCK was inhibited. Since ROCK increases the activity of myosin, this is similar another study were the effect of myosin inhibitors were found to have a stiffening effect on suspended cells ^14^. It was proposed that myosin-induced actin depolymerization played a role. However ROCK also inactivates cofilin, through the activation of LIM-kinase, inhibiting actin depolymerization^53^ making the cause of the observed effect unclear.

Mechanical mismatch between transformed and untransformed cells may be an important factor in metastasis. During early tumor growth cancer cells are in contact with untransformed cells in an epithelial monolayer. It has been shown that the migration speed of single, mechanically soft cells are enhanced by surrounding stiff cells ^54^. The mismatch may also play a role in the extrusion of cancer cells. In healthy epithelia, cells that are destined to die are extruded from the monolayer by the actomyosin contraction of the surrounding cells ^55^. Cells in epithelial monolayers undergoing mitosis also round up through increasing their cortical tension by an increase in actomyosin contractility ^56–58^. Cells extruded from monolayers die by anoikis, the programed cell death of anchorage- dependant cells ^59^. Transformed epithelial cells are also extruded from the monolayer ^60,61^. Normally, epithelia extrude cells apically into the external environment, which would essentially eliminate transformed cells ^62^. However, oncogenic cells can also extrude basally under the epithelium, which could potentially initiate metastasis ^25,62^. Importantly, these processes of extrusion occur only when wild-type and transformed cells contact one another in an epithelial monolayer not when transformed cells are alone ^25,40^. In our experiments, the Ras^V12^ cells were found to form basal protrusions and apically extrude when grown in co-culture with WT cells. Untreated Ras cells demonstrated basal protrusions more often than apical extrusion. However with the addition of the ROCK inhibitor Y-27632, we observed more apical extrusion and less basal protrusion.

Our finding that Ras^V12^ cells were softer than WT cells in co-culture when treated with a ROCK inhibitor may explain this result. The mechanical mismatch, both with the transformed cells being stiffer or softer than the untransformed cells, was accompanied with apical extrusion and basal protrusions from the monolayer. However, when the transformed cells were softer than the surrounding cells, they had more protrusion and less extrusion than when they were stiffer. The monolayer extrusion mechanism may favor basal protrusion for stiffer cells and apical extrusion for softer cells.

The mechanical properties of cells may play a crucial role in cancer progression. After a cell is transformed by a mutation, the cell is either eliminated or it is allowed to proliferate and metastasize by spreading to new sites. The elastic mismatch may play an energetic role in the extrusion of the cells from the monolayer. There may be a link between the softening effect of Y-27632 on Ras^V12^ cells and decreased basal extrusion, suggesting extrusion processes may be dependant on the stiffness of the cells. After the cells enter the blood stream, they must also infiltrate the tissue at the secondary site, where they will form a metastasis ^63^. Circulating tumor cells pass through narrow vessels, smaller than their diameters ^64^. It is believed that increased cell compliance aids in this process. It has been shown that that circulating blood cells differentiate to acquire a compliant phenotype in order to pass through constrictions in blood vessels ^65^. Although neither of attachment states examined in this study is fully equivalent to the situation *in vivo*, it demonstrates that the relative stiffness of oncogenic Ras transformed cells is dependant on the degree of attachment. This may play a distinct role in the metastasis process, where attachment levels vary at each step.

## CONCLUSIONS

Constitutively active Ras has opposite effects on the relative stiffness of attached and unattached epithelial cells. This was likely a result of the Ras-rho-ROCK pathway having opposite effects on suspended and attached cells. Our results also suggest that Ras transformed cells are inherently softer, as shown when the effects of ROCK were inhibited by Y-27632. This may play a role in cancer cell extrusion, protrusion and migration. This information could be used to further understand the effect of Ras on cell mechanics and potentially aid in the development of treatments that target them.

## ACKNOWLEDGMENTS

This work was supported by a Natural Sciences and Engineering Research Council (NSERC) Discovery Grant. C.G. was supported by the uOttawa NSERC-CREATE program in Quantitative Biomedicine. A.E.P. acknowledges support from the Canada Research Chairs (CRC) program. J.G. acknowledges support from the ERC Starting Grant "LightTouch" (grant agreement number 282060).

